# Calcineurin-dependent contributions to fitness in the opportunistic pathogen *Candida glabrata*

**DOI:** 10.1101/2023.09.25.559325

**Authors:** Matthew W. Pavesic, Andrew N. Gale, Timothy J. Nickels, Abigail A. Harrington, Maya Bussey, Kyle W. Cunningham

## Abstract

The protein phosphatase calcineurin is vital for virulence of the opportunistic fungal pathogen *Candida glabrata*. The host-induced stresses that activate calcineurin signaling are unknown, as are the targets of calcineurin relevant to virulence. To potentially shed light on these processes, millions of transposon insertion mutants throughout the genome of *C. glabrata* were profiled *en masse* for fitness defects in the presence of FK506, a specific inhibitor of calcineurin. 87 specific gene deficiencies depended on calcineurin signaling for full viability *in vitro* both in wild type and *pdr1Δ* null strains lacking pleiotropic drug resistance. Three genes involved in cell wall biosynthesis (*FKS1*, *DCW1*, *FLC1*) possess co-essential paralogs whose expression depended on calcineurin and Crz1 in response to micafungin, a clinical antifungal that interferes with cell wall biogenesis. Interestingly, 80% of the FK506-sensitive mutants were deficient in different aspects of vesicular trafficking, such as endocytosis, exocytosis, sorting, and biogenesis of secretory proteins in the ER. In response to the experimental antifungal manogepix that blocks GPI-anchor biosynthesis in the ER, calcineurin signaling increased and strongly prevented cell death independent of Crz1, one of its major targets. Comparisons between manogepix, micafungin, and the ER-stressing tunicamycin reveal a correlation between the degree of calcineurin signaling and the degree of cell survival. These findings suggest that calcineurin plays major roles in mitigating stresses of vesicular trafficking. Such stresses may arise during host infection and in response antifungal therapies.

## INTRODUCTION

Calcineurin is a serine/threonine protein phosphatase that is strongly conserved from fungi to animals whose activity depends on the binding of Ca^2+^ and Ca^2+^/calmodulin (1). Two different natural products, cyclosporin A and FK506, potently bind and inhibit calcineurin signaling in both fungi and animals. These inhibitors have been utilized extensively in clinical settings as immunosuppressants, as the NFAT transcription factors in human T-cells critically depend on calcineurin signaling for triggering normal immune responses (2). These calcineurin inhibitors also impact calcineurin signaling in neurons and other cell types in the heart, kidneys, and muscle probably through a variety of different NFAT-independent processes (3).

Calcineurin inhibitors do not impact the growth of most yeasts in unstressed laboratory conditions. Similarly, deletion of genes encoding the catalytic (*CNA1*) or regulatory (*CNB1*) subunits of calcineurin in yeasts does not impact growth (4–8). However, calcineurin deficiency strongly diminishes virulence of many fungal pathogens in animal models of fungal disease. The pathogenic yeasts *Candida albicans* (9–11), *C. tropicalis* (12)*, C. dubliniensis* (13), and *C. glabrata* (6, 7) strongly depend on calcineurin function for colonization, proliferation, or survival in mouse models of invasive candidiasis. The mold *Aspergillus fumigatus* (14, 15) and the basidiomycete *Cryptococcus neoformans* (16) also depend on calcineurin for successful colonization and disease progression. During host infections, pathogenic fungi may experience stresses that trigger calcineurin activation and dephosphorylation of specific targets involved in mitigating those stresses or prolonging cell survival.

Calcineurin also promotes resistance and tolerance in pathogenic fungi to several different classes of antifungals (17, 18). For example, calcineurin activation and signaling promotes tolerance (also called cell viability) during long-term exposure of yeasts to azole-class antifungals, which target ergosterol biosynthesis in the endoplasmic reticulum (ER) (19–21). Similar effects have been observed in response to non-clinical antifungals such as tunicamycin and dithiothreitol, which perturb glycoprotein biosynthesis in the ER (20, 22). The pro-survival effects of calcineurin during ER stress can operate independent of Crz1, a fungi-specific transcription factor unrelated to NFAT that is directly dephosphorylated by calcineurin (20, 22). Several other known substrates of calcineurin in the model yeast *S. cerevisiae* were also not required for calcineurin-dependent cell survival in response to ER stress (23) and the molecular mechanisms by which calcineurin promotes tolerance remain unknown. Calcineurin and Crz1 activation also promote echinocandin resistance by increasing expression of *FKS2* encoding a target of these drugs (17, 18). Echinocandin resistance often arises through mutations within the coding sequences of *FKS* genes (24, 25), although evidence suggests other pathways can contribute (26). Based on these findings, selective inhibitors of fungal calcineurin would enhance the potency or cidality of existing antifungals and diminish intrinsic virulence of fungal pathogens without causing immunosuppression in the host (27).

A better understanding of the stresses in fungi that are sensed by calcineurin and how the sensor promotes resistance, tolerance, and virulence could provide new approaches for antifungal interventions. Toward that goal, this study aims to identify cellular stresses in *C. glabrata* where calcineurin signaling becomes essential for fitness *in vitro*. These *in vitro* stresses may be representative of stresses encountered *in vivo* during host infections, thereby facilitating deeper studies of the inputs and outputs of the calcineurin signaling pathway. By profiling large pools of transposon insertion mutants in *C. glabrata* for hypersensitivity to FK506, a broad range of genotoxic stresses can be surveyed. Using this approach, we identify dozens of genes and several cellular processes whose disruptions cause dependence on calcineurin signaling for fitness. One such process is cell wall biosynthesis, in which calcineurin appears to induce paralogs that function in compensatory pathways and provide resistance to an echinocandin. GPI-anchor deficiency, which is also conferred by the experimental antifungal manogepix, also produces a cellular stress for which calcineurin function compensates. Overall, deficiencies in vesicular trafficking were the most common cellular stresses that depended on calcineurin signaling. These genetic stresses *in vitro* may model the physiological stresses encountered by *C. glabrata* during infections of animal hosts.

## RESULTS

### Mutants with elevated dependence on calcineurin signaling

We sought to identify stresses in *C. glabrata* during which calcineurin becomes important for cell survival or proliferation. To do this, we cultivated complex pools of *Hermes* transposon insertion mutants in the strain BG14 (28), a *ura3Δ* derivative of vaginal isolate BG2 (29), in the presence and absence of FK506 for 6.7 generations. Each insertion site was PCR amplified, sequenced, mapped, and tabulated gene-wise as described previously (30), and then each gene was charted under the two conditions (Fig. 1A). Most genes were equally represented by transposon reads in the two conditions, suggesting their disruptions do not generate stresses that are compensated by calcineurin signaling (black dots in Fig. 1A). No disrupted genes were significantly enriched in the FK506 condition. However, dozens of disrupted genes were significantly underrepresented following FK506 exposure. Among those genes is the known FK506-sensitive gene *FKS1* (red square) that was shown previously to depend on calcineurin signaling when disrupted (31). The experiment was repeated in an independently generated pool of insertion mutants in the BG14 background and also in an independently generated pool of insertion mutants in a *pdr1Δ* derivative of BG14 (32). This *pdr1Δ* pool was profiled because FK506 may regulate Pdr1 transcription factor independent of calcineurin (33) and because calcineurin can physically interact with Pdr1 (34). For each experiment, z-scores were calculated for all annotated genes and compared. *FKS1* exhibited strongly negative z-scores in all three experiments (−8.9, −17.2, −9.6). Overall, the z-scores from the two BG14 experiments were highly correlated with each other (PCC = 0.41) and both correlated well with z-scores from the *pdr1Δ* experiment (PCC = 0.36 and 0.47). When the z-scores from *pdr1Δ* experiments were charted against the average z-scores of the replicate BG14 experiments, few outliers were evident (Fig. 1B). These findings suggest dozens of genes reproducibly impact FK506 sensitivity in these conditions independent of Pdr1.

**Figure 1.**
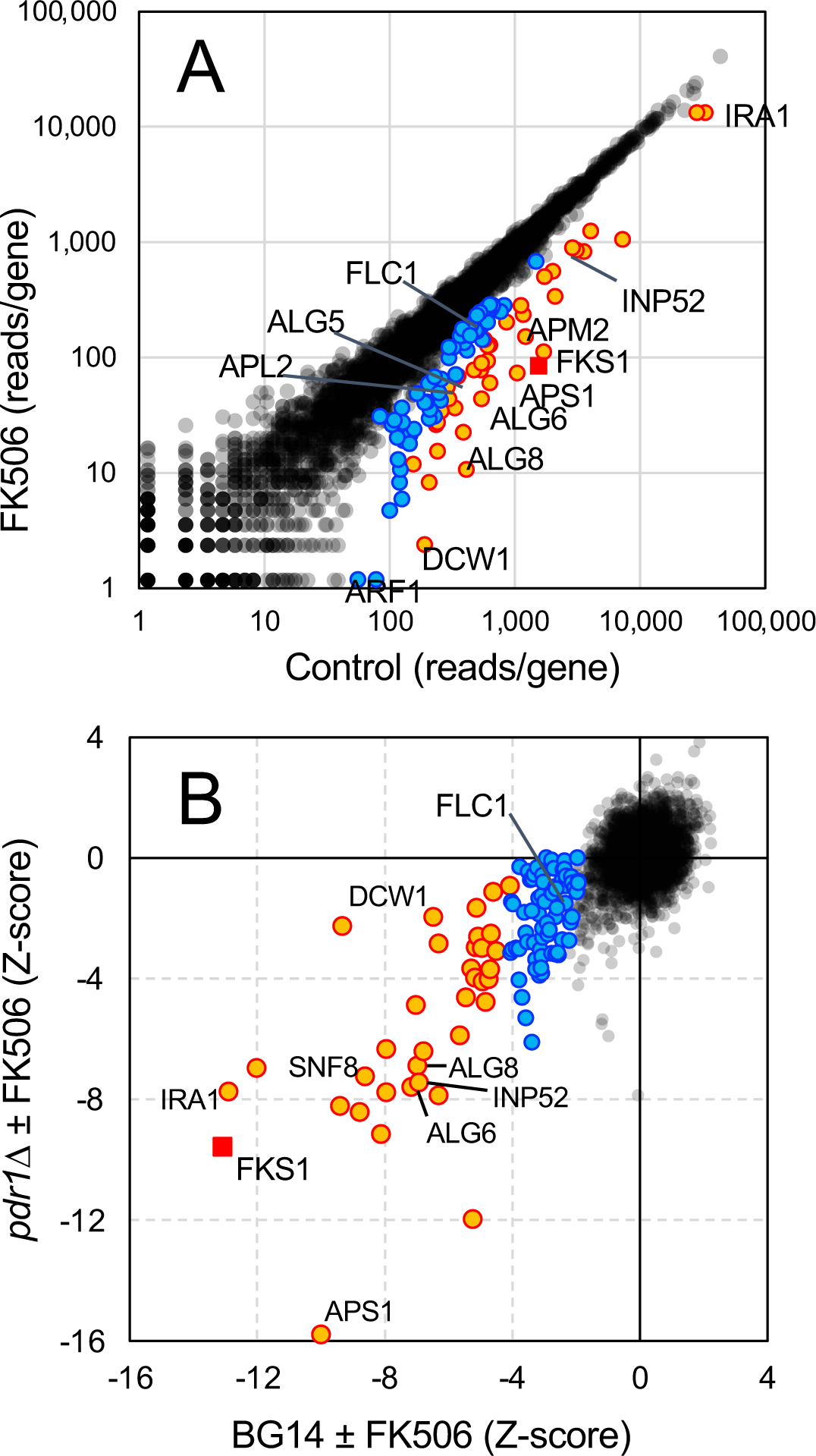
Genome-wide analyses of FK506 susceptibility in *C. glabrata*. (A) A diverse pool of *Hermes* transposon insertion mutants in the BG14 strain background was diluted 100-fold into fresh SCD medium containing or lacking FK506 (1 µg/mL), shaken for one day, and then the insertion sites were sequenced and tabulated gene-wise. Genes falling 2 to 4 (blue) and greater than 4 (orange) standard deviations below the main diagonal are highlighted. (B) z-scores from panel A and an independent pool of insertion mutants in BG14 were averaged and plotted against Z-scores obtained similarly from a pool of insertion mutants in a *pdr1Δ* derivative of BG14.

After averaging all three datasets, a total of 87 genes were depleted more than 2-fold and with z-scores less than −2.0 in response to FK506 (Table S1). The *SSD1* gene was not significant in these datasets even though *ssd1Δ* mutants were shown previously to be hypersensitive to FK506 (35). Closer inspection of *SSD1* revealed the gene was poorly covered with transposon insertions, suggesting it was essential for viability in these conditions. Gene Ontology analyses revealed >4-fold enrichment of ten processes, nine of which involve vesicular trafficking (Table S2). When z-scores were limited to less than −4.0 (34 genes), six processes were enriched >4-fold and all involved vesicular trafficking. After manual curation, over 79% of the significant genes were found to contribute in some way to vesicular trafficking with the remainder contributing to cell wall biogenesis or unidentified functions.

### Co-essential paralogs that depend on Crz1 and/or calcineurin

The *FKS1* gene encodes a catalytic subunit of beta-1,3-glucan synthase, which is also the molecular target of micafungin and other echinocandin-class antifungals. *FKS1*-deficient mutants of *C. glabrata* are viable but hypersensitive to FK506 because expression of *FKS2*, a functionally redundant co-essential paralog, depends on calcineurin and Crz1 (6, 31). To determine whether other FK506-hypersensitive mutants might depend on calcineurin and Crz1 for expression of co-essential paralogs, we scanned the top 87 genes for paralogs and then searched published genetic interaction datasets from *S. cerevisiae* for evidence of co-essentiality (36). Of the 87 genes found in our screen, 15 genes have paralogs in *C. glabrata*, 9 of which are conserved and co-essential in *S. cerevisiae* (Table S1). We then quantified expression patterns of all 9 co-essential paralogs relative to control genes using RT-PCR in wild-type and *crz1Δ* mutant cells with and without exposure to micafungin (see methods). These conditions activated calcineurin and induced expression of *RCN2,* a well-studied target of calcineurin and Crz1 in *C. glabrata* (7, 37), in a Crz1-dependent fashion (Fig 2). The same pattern of expression was observed for *FKS2*, as expected from earlier studies (6, 31). Five co-essential paralogs (*ARF2*, *INP53*, *IRA2*, *PMT3*, *SEC12*) and seven other non-essential paralogs (*BOI2*, *CCW12*, *FKS3*, *PEX31*, *UPC2*, *VPS501*, *YEH1*) exhibited no significant response to micafungin and/or Crz1 deficiency (Fig. 2). One co-essential paralog (*MYO3*) responded to micafungin independent of Crz1 and was not further studied. Interestingly, two co-essential paralogs of *DCW1* and *FLC1* (*DFG5* and *FLC2*) were expressed similar to *FKS2* and *RCN2* (Fig. 2), suggesting co-regulation by calcineurin and Crz1. Previous studies in *S. cerevisiae* (38) showed that calcineurin and Crz1 can induce expression of *DFG5*, *FLC2*, *FKS2*, and *RCN2* orthologs, indicating evolutionary conservation of the regulatory network.

**Figure 2.**
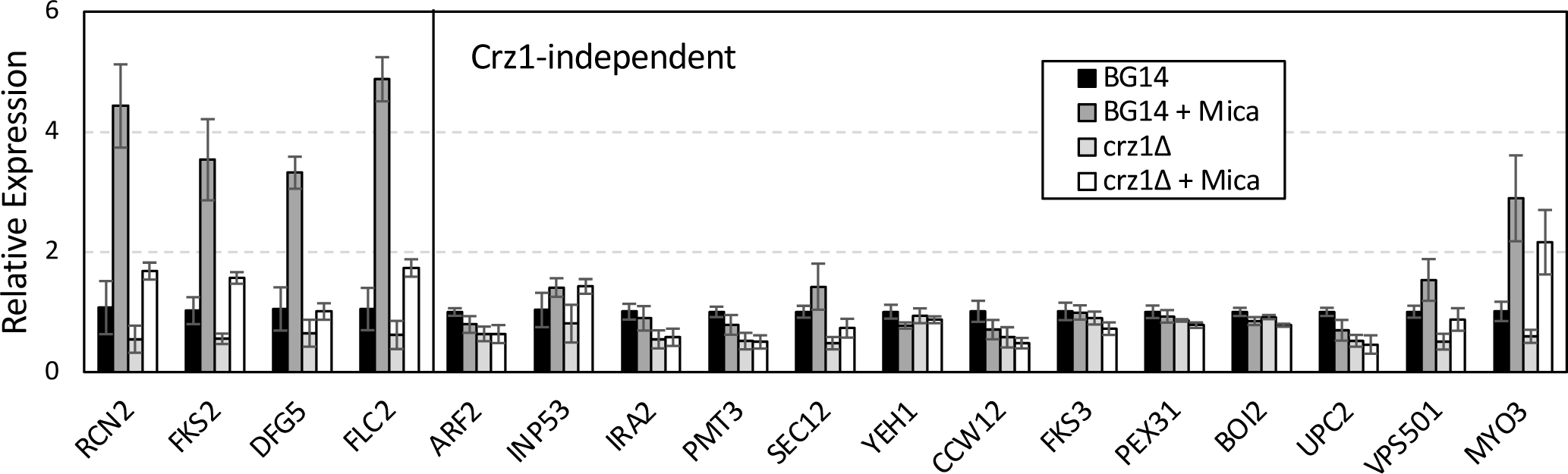
Calcineurin and Crz1 regulate expression of *FKS2*, *DFG5*, and *FLC2* but not other co-essential paralogs in response to cell wall stressor (micafungin). Expression of the indicated genes was monitored by qRT-PCR in wild-type BG14 cells and a *crz1Δ* derivative with and without exposure to micafungin for 80 minutes. Columns indicate averages of 4 replicates (±SD).

### Crz1-dependent co-essential paralogs in the cell wall

*FKS1* and *FKS2* encode transmembrane enzymes that synthesize beta-glucan in the cell wall. *DCW1* and *DFG5* encode enzymes that are GPI-anchored to the plasma membrane (39) and catalyze the covalent attachment of numerous other GPI-anchored proteins, such as adhesins, to the beta-glucan in the cell wall (40, 41). *FLC1* and *FLC2* encode transmembrane proteins of the ER with incompletely established functions in *S. cerevisiae* (42, 43). Through analysis of published datasets, we obtained evidence suggesting that *FLC1* and *DCW1* gene products form a functional partnership and that *FLC2* and *DFG5* gene products form a similar redundant partnership. First, the products of *FLC2* and *DFG5* physically interacted in *S. cerevisiae* (44). Second, *flc2Δ* and *dfg5Δ* mutants exhibited strikingly similar chemical interaction profiles (45) and cluster together when analyzed alongside thousands of other knockout mutants (46). Third, strong fitness defects (synthetic lethalities) were observed in double mutants lacking both *DCW1* and *FLC2* and both *DCW1* and *DFG5*, while no fitness defects were observed in double mutants lacking both *DCW1* and *FLC1* or *DFG5* and *FLC2* (36) as illustrated (Fig. 3A). All these findings support a hypothesis that *DFG5* and *FLC2* function together in the same complex or pathway that is functionally redundant with the *DCW1* and *FLC1* complex or pathway in *S. cerevisiae*.

**Figure 3.**
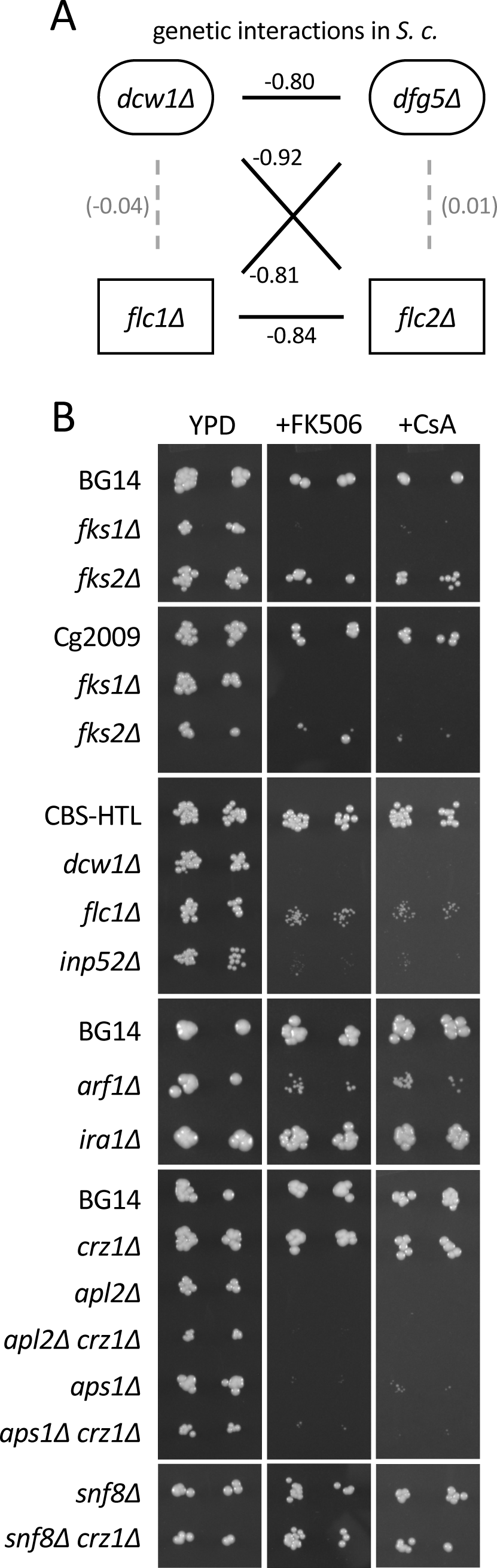
Responses of individual knockout mutants of *C. glabrata* to calcineurin inhibitors. (A) Genetic interactions of gene knockout mutants in *S. cerevisiae* obtained from (36). (B) Serial dilutions of the indicated *C. glabrata* strains were frogged onto SCD agar medium containing calcineurin inhibitors, incubated for 1 day at 30°C, and photographed.

Deficiencies in *FKS1*, *DCW1*, or *FLC1* may cause cell wall stresses in *C. glabrata* that activate calcineurin and Crz1, which could increase expression of *FKS2*, *DFG5*, *FLC2* paralogs in a compensatory fashion. To further explore this hypothesis, *fks1Δ*, *dcw1Δ*, and *flc1Δ* single knockout mutants of *C. glabrata* were studied. In both the BG2 and Cg2009 strain backgrounds, the *fks1Δ* mutants grew very poorly in medium supplemented with FK506 or cylcosporin A, an alternative inhibitor of calcineurin (Fig. 3B), as expected from earlier studies. The *dcw1Δ* and *flc1Δ* mutants also grew poorly in the presence of both calcineurin inhibitors relative to the control strain CBS138-HTL (47) (Fig. 3B). To determine whether calcineurin and Crz1 were activated in these mutants, expression of *RCN2* was monitored by RT-PCR. Basal expression of *RCN2* was significantly elevated in *fks1Δ*, *dcw1Δ*, and *dfg5Δ* mutants relative to the control strains, and this effect was blocked by FK506 in most cases (Fig. 4). Thus, the genetic losses of *FKS1*, *DCW1*, and *FLC1* seemed to cause stresses that activated calcineurin and Crz1 similar to micafungin exposure, which in turn increased expression of the co-essential paralogs that serve to bolster cell wall biosynthesis and remodeling.

**Figure 4.**
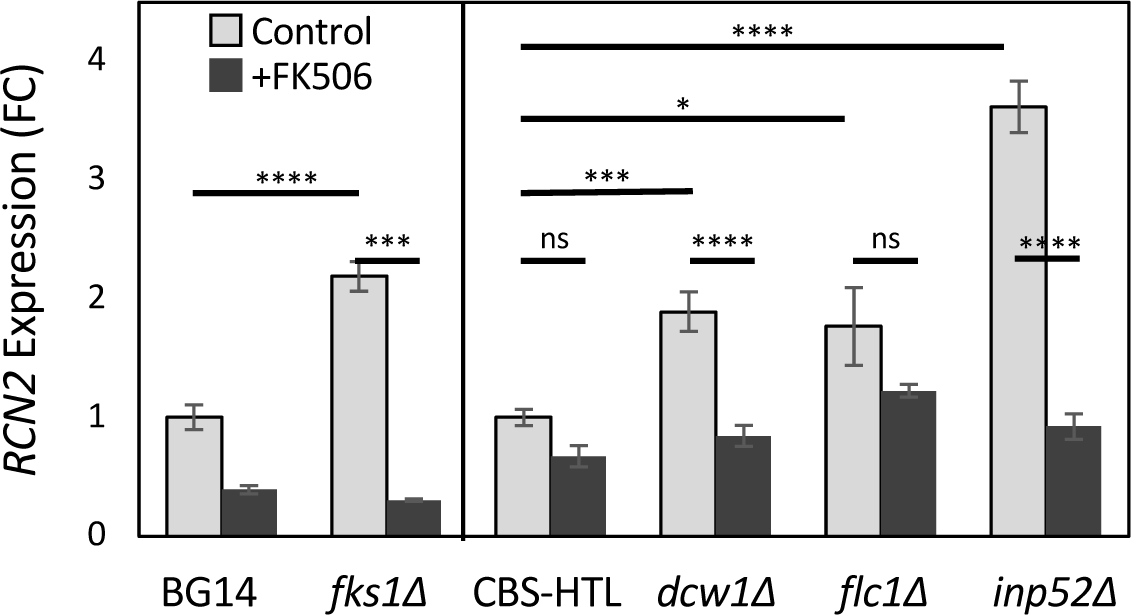
Genetic deficiencies in cell wall biogenesis increase calcineurin signaling. Expression of the calcineurin- and Crz1-dependent gene *RCN2* was monitored by qRT-PCR in the indicated strains during exponential growth in SCD medium at 30°C with or without exposure to FK506. Columns indicate averages of 4 replicates (±SD). Statistical significance was assessed using a Welch’s T-test with Bonferroni correction for multiple comparisons (*, P < 0.05; **, P < 0.01; ***, P < 0.005; ****, P < 0.001; ns, not significant).

### *YPS7*-deficient mutants depend on calcineurin

Transposon insertions suggest *YPS7* deficiency alone causes hypersensitivity to FK506 (average Z-score = −2.9). The *YPS7* gene encodes one of eleven cell wall anchored and secreted aspartyl proteases (SAPs, or yapsins) that contribute to cell wall remodeling but are not co-essential *in vitro* (48). Yapsin deficiencies diminished shedding of Epa1 and other adhesins that are anchored to the cell wall by the actions of Dcw1 and Dfg5 (41, 48). Two other yapsins (*YPS1*, *YPS5*) have been shown to depend on Crz1 and calcineurin for maximum expression (7, 49). In drop tests, we confirmed that *yps7Δ* knockout mutants in the BG14 strain background exhibited strong hypersensitivity to FK506 (Fig. 5). Similar results were obtained in a *yps1Δ* strain background and a *yps5cΔ* background that also lacked *YPS2* and a cluster of eight *YPS* genes including *YPS5* (Fig. 5). A strain bearing only *YPS7* and lacking ten other yapsins retained an ability to grow in FK506, while the strain lacking all eleven yapsins exhibited FK506 hypersensitivity (Fig. 5). These findings suggest that calcineurin promotes growth or survival of *yps7Δ* cells through compensatory effects on non-yapsin targets that remain to be identified. This compensatory effect of calcineurin could be important for virulence, as yapsins already have been shown to be required for virulence of *C. glabrata* in mouse models of invasive candidiasis (48).

**Figure 5.**
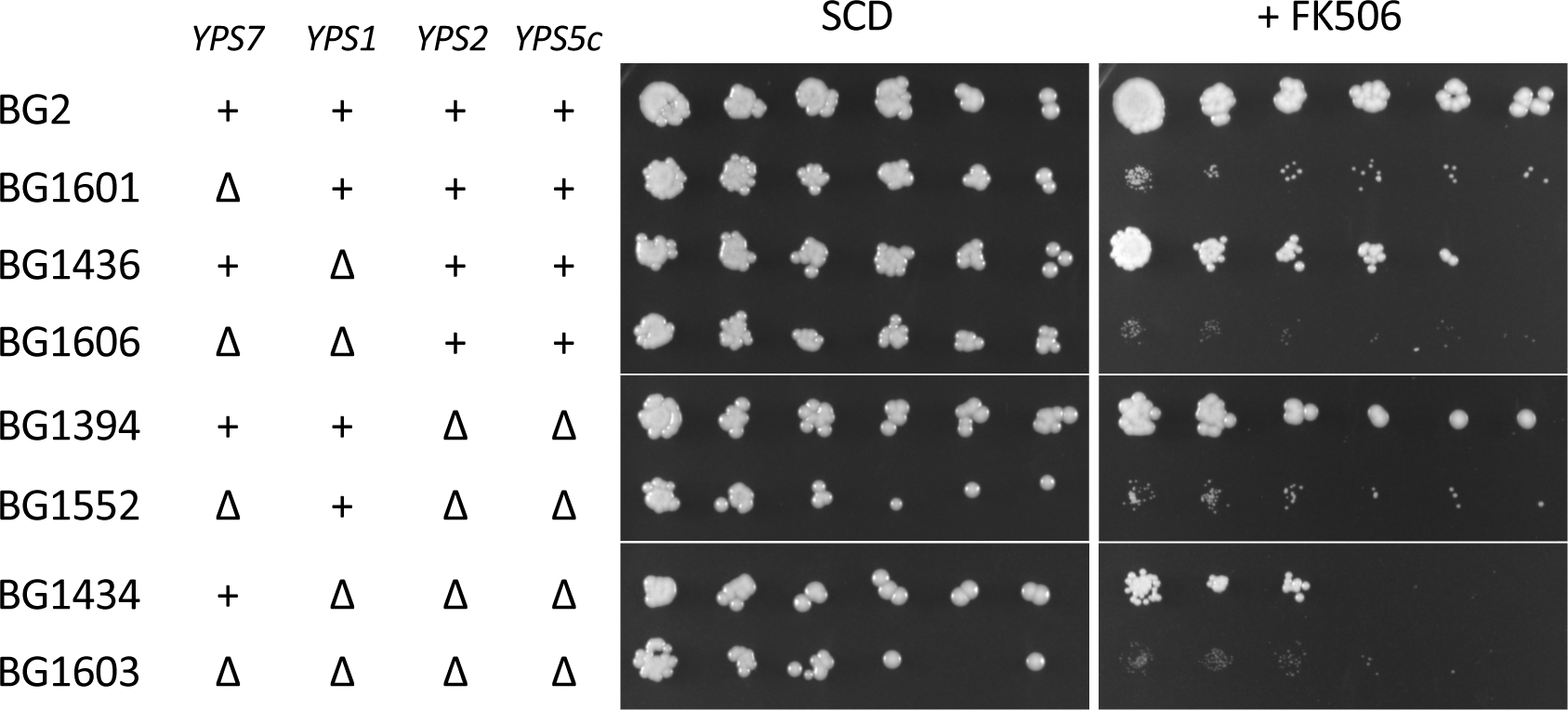
Calcineurin increases fitness of *yps7Δ* mutants independent of ten yapsins. The indicated strains were serially diluted and frogged onto SCD agar medium with or without FK506 as described in Fig. 3. The slower growing BG1434 and BG1603 strains were photographed after 2 days of incubation at 30°C while the others were photographed after 1 day.

### Crz1-independent Co-essential paralogs

Of the five genes whose co-essential paralogs were not responsive to micafungin or Crz1 (Fig. 2), cells lacking either *ARF1* or *INP52* were found to be highly sensitive to FK506 and cyclosporin A (Fig. 3B). Two other strains (lacking either *IRA1* or *PMT2*) were insensitive to the calcineurin inhibitors (Fig. 3B; data not shown). These findings suggest calcineurin may become activated in *arf1Δ* and *inp52Δ* mutants and compensate for the deficiencies through Crz1-independent effects. Consistent with this idea, *RCN2* expression was found to be highly elevated in *inp52Δ* cells relative to wild-type and lowered upon exposure to FK506 (Fig. 4). Calcineurin can directly dephosphorylate and regulate the *INP53* gene product in *S. cerevisiae* (50). These findings suggest that calcineurin may compensate for some types of cellular stresses through Crz1-independent processes.

### Calcineurin improves fitness during stresses in vesicular trafficking

*INP52, ARF1,* and their paralogs encode proteins involved in different aspects of vesicular trafficking, such as endocytosis and the formation of coated vesicles. Strikingly, defects in vesicular trafficking and endoplasmic reticulum functions were predicted for 69 of the 87 different mutants identified above as FK506-hypersensitive, of which 67 do not have identifiable paralogs (Table S1). The mutant genes include *VPS13*, *GGA1* and *CHS6* whose products interact with *ARF1* in *S. cerevisiae* (tabulated at yeastgenome.org (51)). The screens found that all of the core subunits of the AP-1 and AP-1R complexes (*APM1, APM2*, *APL2*, *APL4*, *APS1*, *LAA1, MIL1*) that bind clathrin and Arf1 and promote vesicular transport from the Golgi complex to endosomes were among this group. Furthermore, we found that multiple subunits of the ESCRT complexes that promote endosomal trafficking, multivesicular body formation, and vacuolar delivery were also FK506-sensitive. We knocked out two genes encoding AP-1 subunits (*APL2*, *APS1*) and one subunit of the ESCRT-II complex (*SNF8*) in the BG14 strain of *C. glabrata* and tested for calcineurin dependence. The *apl2Δ* and *aps1Δ* mutants were strongly hypersensitive to the calcineurin inhibitors, though both formed slightly smaller colonies than the parent strain in the absence of calcineurin inhibitors (Fig. 3B). The *snf8Δ* mutant exhibited very weak sensitivity to calcineurin inhibitors in these conditions (Fig. 3B), which contrasts with the large effects observed in the transposon pools (average z-score = −8.2). *SNF8* is a large gene with high density of transposon insertions in our pools, which causes even small phenotypic effects to be highly significant in our z-score calculations. The three genes were also knocked out in a *crz1Δ* mutant background and tested similarly. The resulting double mutants behaved indistinguishable from the *apl2Δ*, *aps1Δ*, and *snf8Δ* single mutants (Fig. 3B), suggesting that calcineurin promotes fitness during these stresses through Crz1-independent effects. The relevant targets of calcineurin have not yet been identified.

### ER stresses activate calcineurin, which promotes growth and cell survival

Several genes whose products promote secretory protein modifications in the ER were on the list of FK506-sensitive mutants with vesicular trafficking defects. Among these were *ALG3*, *ALG5*, *ALG6*, and *ALG8* whose non-essential products function sequentially in the N-glycan biosynthetic pathway, *CNE1* that binds glucosylated N-glycosylated secretory proteins, and *GTB1* that promotes their deglucosylation. A *cne1Δ* mutant of *C. glabrata* was previously shown to be hypersensitive to FK506 (52). The *alg5Δ*, *alg6Δ*, and *alg8Δ* mutants of the BG14 background all exhibited mild sensitivity to calcineurin inhibitors (Fig. 6A). Elevated levels of *RCN2* expression were observed in *alg6Δ* and *alg8Δ* mutants (Fig. 6B). Additionally, several genes involved in GPI-anchor biosynthesis in the ER (*LAS21*, *PER1*, *BST1*) and trafficking of GPI anchored glycoproteins from the ER to the Golgi complex (*EMP24*) were also identified as hypersensitive to FK506 when disrupted with transposons. A *las21Δ* mutant in the CBS138-HTL background exhibited hypersensitivity to FK506 and cyclosporin A (data not shown). Insertions in the 354 bp gene *CAGL0G03993g* were highly significant in all three pools (average z-score = − 5.2). This small gene is not transcribed (53), and its product is not conserved in any other species, suggesting it is misannotated as a gene. Furthermore, the segment begins only 38 bp upstream of the essential gene *GPI13*, which encodes an enzyme critical for GPI-anchor biosynthesis in the ER. Therefore, insertions in the *CAGL0G03993g* segment likely lower the expression of *GPI13*, potentially causing stresses similar to insertions in the non-essential GPI-anchor genes.

**Figure 6.**
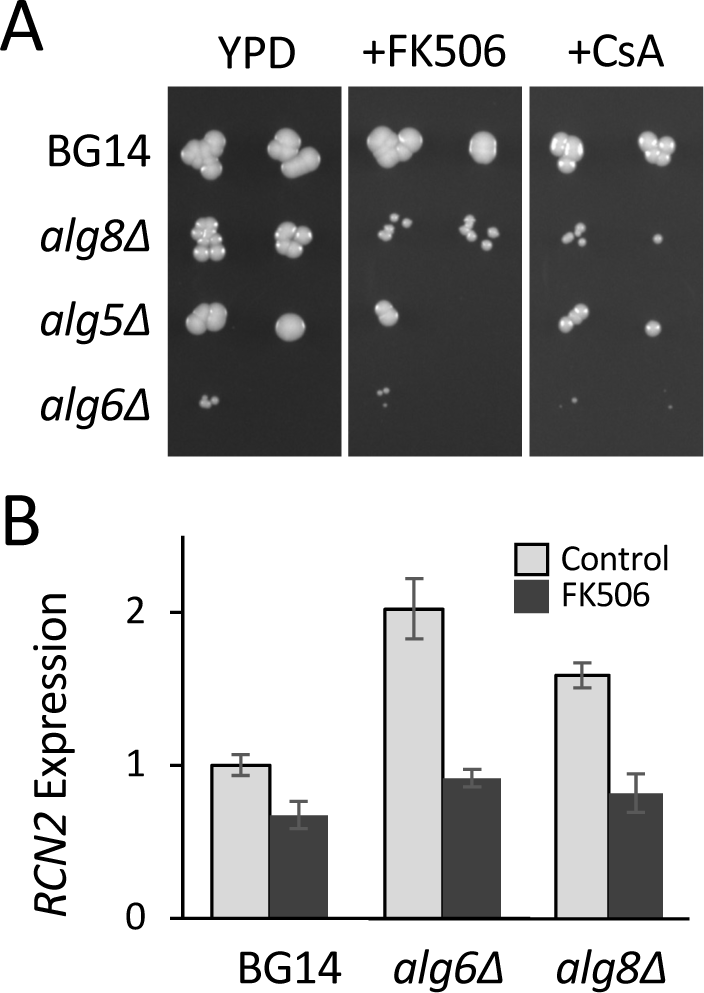
Genetic deficiencies of N-glycosylation in the ER increase calcineurin signaling and dependence. Drop tests (A) and qRT-PCR measurements (B) were performed on the indicated strains as described in Figures 3 and 4.

Essential genes are poorly represented with transposon insertions, which limits their detectability in the genetic screens. To determine whether calcineurin can become activated in response to deficiencies in essential components of N-glycosylation and GPI-anchoring, we quantified *RCN2* expression after exposure of wild-type and *crz1Δ* mutant cells to tunicamycin and manogepix. Manogepix is an experimental antifungal that blocks an intermediate step in GPI-anchor biosynthesis in the ER encoded by essential *GWT1* (54). Tunicamycin is a natural product that blocks the first step of N-glycan biosynthesis in the ER encoded by essential *ALG7* (55). Addition of manogepix and tunicamycin to exponentially growing cells at high concentrations caused rapid induction of *RCN2* expression in wild-type cells but not in *crz1Δ* mutant cells (Fig. 7A). Interestingly, the degree of *RCN2* induction was much higher for these ER stressors than for micafungin (Fig. 7A). These findings suggest that both inhibitors produce cellular stresses that rapidly and strongly activate calcineurin signaling.

**Figure 7.**
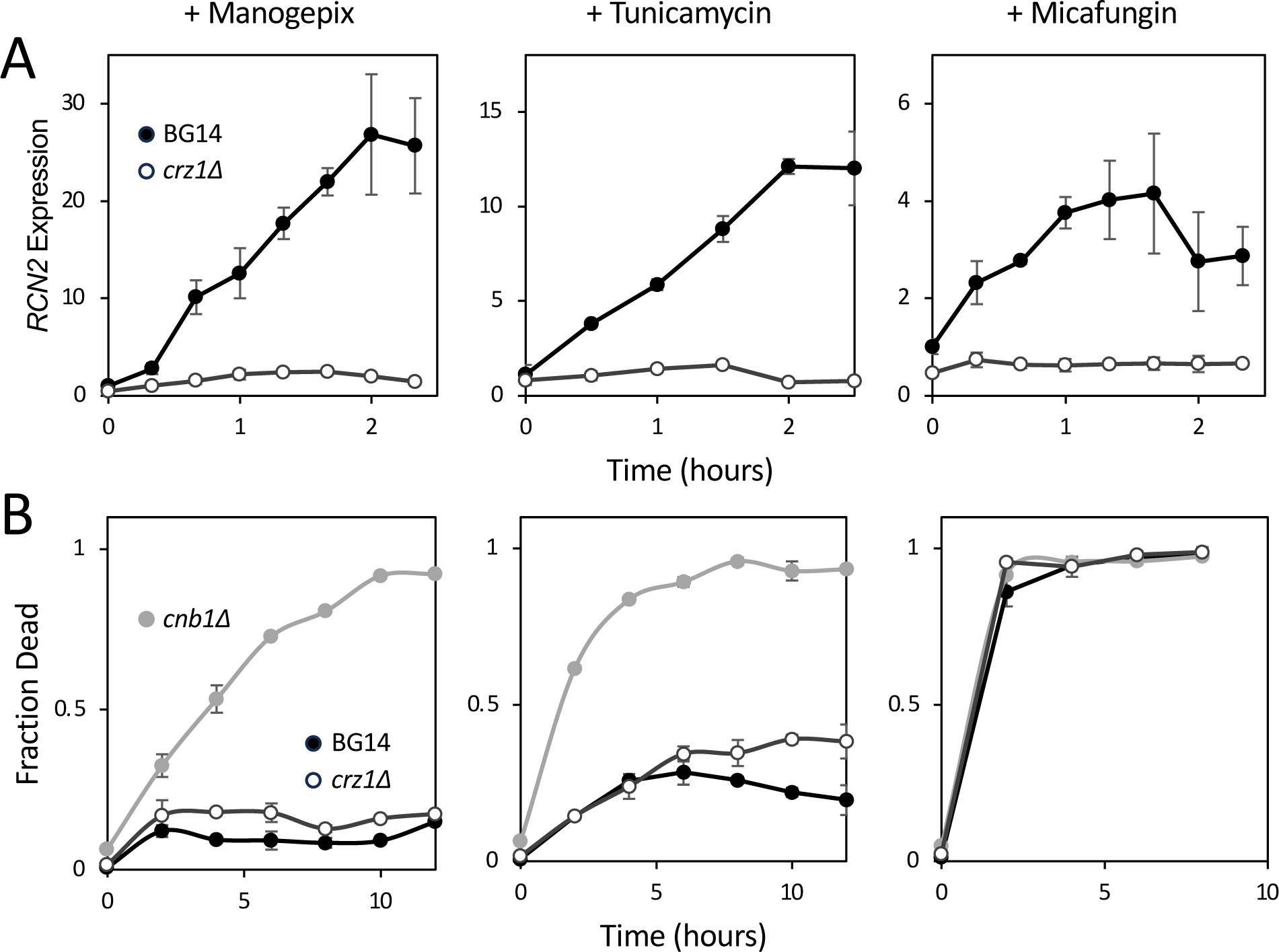
Acute inhibition of N-glycosylation and GPI-anchoring in the ER trigger calcineurin signaling which promotes cell survival. BG14 (black symbols), *crz1Δ* (white symbols), and *cnb1Δ* (gray symbols) cells were grown to log phase in SCD medium at 30°C and then exposed to manogepix (0.6 µg/mL), tunicamycin (20 µg/mL), or micafungin (0.12 µg/mL). At the indicated times, samples were removed and analyzed by qRT-PCR to measure *RCN2* expression and by propidium iodide staining to measure cell death in the population (B). Data points indicate the averages of 3 biological replicates (±SD).

In *S. cerevisiae,* calcineurin promotes cell survival during exposure to tunicamycin independent of Crz1 (20). To test whether calcineurin performs similar roles in *C. glabrata*, *crz1Δ* and *cnb1Δ* mutants were exposed to tunicamycin, and cell death was quantified by staining with propidium iodide. The *crz1Δ* mutant and BG14 control cells largely survived the stresses caused by tunicamycin, whereas the *cnb1Δ* mutant rapidly died (Fig. 7B). Manogepix produced similar effects, except the rate of *cnb1Δ* mutant cell death appeared somewhat slower than that of tunicamycin (Fig. 7B). In contrast, micafungin exposure caused very rapid cell death in all three *C. glabrata* strains (Fig. 7B). For these three antifungals, the rates of cell death in *cnb1Δ* cultures were inversely correlated with the magnitudes of calcineurin activation in wild-type cultures. This correlation between longer cell survival and higher calcineurin signaling is consistent with calcineurin as a driving factor in antifungal tolerance.

## DISCUSSION

This study explores the genetic stresses under which calcineurin signaling becomes important for growth and survival of *C. glabrata* in laboratory conditions. Such *in vitro* stresses may model stresses encountered by the pathogen during host infections, thereby providing insights into how calcineurin signaling may promote virulence. Strikingly, nearly 80% of the genetic stresses where calcineurin became crucial for fitness involved deficiencies in the endoplasmic reticulum and vesicular trafficking systems. Overall, these findings of FK506 sensitivity in *C. glabrata* appear more similar to the findings in the evolutionarily distant fission yeast *Schizosaccharomyces pombe* (56) than to those in the much closer relative *Saccharomyces cerevisiae* (57–59). Some of the differences may be attributed to very different experimental methods. The previous studies utilized large collections of gene knockout mutants that were tested individually. In contrast, this study employs large pools of transposon insertion mutants that were competing against one another and profiled *en masse* by deep sequencing. This method yielded a z-score for each gene. The z-score is sensitive to both the magnitude of the phenotype and the intrinsic noise, which can be very large for small genes and essential genes that contain few transposon insertions relative to other genes in the pool. For large transposon-rich genes such as *IRA1*, highly significant z-scores can be obtained with low effect size. Conversely, insignificant z-scores could arise for genes with high phenotypic sensitivity to FK506 but very low transposon coverage. This effect may explain why *SSD1* was not identified in our datasets in spite of its previously established hypersensitivity to FK506 in a different *C. glabrata* strain (35). When *S. cerevisiae* gene knockouts were screened individually for elevated Ca^2+^ uptake and calcineurin signaling rather than FK506 sensitivity (60), numerous genes involved in ER and vesicular trafficking processes were identified. An emerging theme from all these studies is that calcineurin activation and signaling plays a broadly conserved role in the compensatory responses to stresses in vesicular trafficking.

The ER stressor tunicamycin has been shown previously to trigger calcineurin dependency and FK506 hypersensitivity in *C. glabrata* as well as *C. albicans* and *S. cerevisiae* (20). The genetic screens performed here reveal several additional non-essential genes of the N-glycosylation process in the ER as well as several non-essential genes of the GPI-anchoring process in the ER that modifies about 135 secretory proteins including the yapsins, adhesins, Dfg5, Dcw1, and others (61). Exploiting manogepix as an acute inhibitor of GPI-anchoring, we showed that calcineurin became activated rapidly, and this activation was critical for *C. glabrata* cell survival independent of Crz1. *C. albicans* also may rely on calcineurin for survival in response to manogepix and other GPI-anchoring deficiencies (62). The pro-survival effects of calcineurin during these forms of ER stress may overlap with the pro-survival effects of calcineurin in yeasts exposed to clinical azoles, which inhibit the *ERG11* gene product required for ergosterol biosynthesis in the ER (63). If the pro-survival substrates of calcineurin can be identified, alternatives to FK506 that promote conversion of these fungistats into fungicides may become possible.

Genetic deficiencies in most major steps of the vesicular trafficking network also resulted in calcineurin dependence in *C. glabrata*. Assuming the genes function similar to orthologs in *S. cerevisiae*, these steps include packaging of N-glycosylated and GPI-anchored proteins in the ER, their trafficking to and further modification in the Golgi complex, and subsequent sorting and trafficking to the plasma membrane. Mutants deficient in endocytosis and trafficking to the vacuole also demonstrated calcineurin dependence. Dozens of additional essential and non-essential gene products accomplish all these processes. In *S. cerevisiae*, several proteins involved in vesicular trafficking have been identified as direct substrates of calcineurin (3, 64). While orthologs of these substrates are conserved in *C. glabrata*, the motifs required for recognition by calcineurin often are not conserved (64). Some calcineurin substrates, such as the products of *LAC1* and *LAG1*, completely lack the canonical docking motifs for calcineurin (65). These genes encode redundant ceramide synthases in the ER and are excellent candidates for involvement in pro-survival functions of calcineurin. Ceramide accumulation is highly toxic to fungal cells, and its detoxification depends on effective transport to the Golgi complex and enzymatic conversion to sphingolipids (66, 67). Calcineurin signaling can inhibit ceramide biosynthesis observed during ER stress (68) and potentially mitigate toxicity when vesicular trafficking has been stressed/impaired. More research will be necessary to test this hypothesis and others in order to determine how calcineurin promotes fitness of *C. glabrata* cells experiencing stresses in the ER and vesicular trafficking system.

Deficiencies in several cell wall biogenesis genes (*FKS1*, *DCW1*, *FLC1*, *YPS7*, and *CCW22*) were strongly dependent on calcineurin for proliferation in our screens. Unlike the gene deficiencies that stress the ER and vesicular trafficking systems, all these cell wall genes have paralogs or homologs in *C. glabrata*, many of which (*FKS2*, *DFG5*, *FLC2*, *YPS1*, *YPS5*) require calcineurin and Crz1 for maximal expression. Although its expression in *C. albicans* was dependent on calcineurin and Crz1 (69), *CCW12* did not appear to be inducible by micafungin in a calcineurin-dependent manner in *C. glabrata*. Simultaneous deletion of *YPS7* and all ten of its paralogs was not lethal in *C. glabrata* (48), and the resulting undecuple mutant lacking all 11 yapsins still exhibited hypersensitivity to FK506. This finding suggests that calcineurin compensated for *YPS7* and general yapsin deficiencies through some other mechanism. In *S. cerevisiae*, *yps7Δ* mutants depended on *CNB1* for fitness but not *CRZ1* (36), suggesting compensatory effects of calcineurin independent of Crz1 in that species. Crz1 was clearly important for expression of the paralogs of *FKS1*, *DCW1*, and *FLC1* and those paralogs exhibited synthetic lethality when disrupted in *S. cerevisiae*. Some unexpected synthetic lethal interactions were also observed, such as *dcw1Δ flc2Δ* and *flc1Δ dfg5Δ* (36). These findings implicate *FLC1* and *FLC2* in cell wall biogenesis as exclusive partners of *DCW1* and *DFG5*, respectively. Supporting this idea, knockout mutants of *FLC2* and *DFG5* exhibited highly similar chemical interaction profiles (45), and the gene products exhibit physical interactions (44) in *S. cerevisiae*. Though knockout mutants of *FLC1* and *FLC2* exhibit cell wall deficiencies in *S. cerevisiae*, the products localized to ER and were necessary for FAD import, which does not have any obvious role in cell wall biogenesis (42). These findings suggest that calcineurin and Crz1 play major roles in the expression of “reserve” cell wall biogenesis genes in response to deficiencies or inhibition of the primary paralogs.

Inhibitors of *FKS1* and *FKS2* gene products (echinocandins) are utilized clinically for treatment of diverse fungal diseases. The *DCW1* and *DFG5* gene products represent excellent targets for development of novel antifungals due to their broad conservation in fungi as well as the extracellular location of their active sites (40). Chemical-genetic screening has revealed at least two compounds that may inhibit *DCW1* (or *FLC1*) based on the high susceptibility of both *dfg5Δ* and *flc2Δ* mutants of *S. cerevisiae* (45). Our findings predict that calcineurin signaling will promote resistance to such inhibitors by up-regulating expression of the targets and/or paralogs, similar to the action of calcineurin on micafungin resistance. Non-immunosuppressive compounds that specifically block fungal calcineurin would likely augment the potency of those antifungals while suppressing overall virulence even in their absence (17). Targeting the factors upstream and downstream of calcineurin that promote cell survival and proliferation also could be effective at controlling fungal infections. This study provides new insights into those factors in *C. glabrata*.

## METHODS

### Strains and culture conditions

A complete list of *C. glabrata* strains used in this study is shown in Table S3. For individual gene knockouts, the coding sequences between start and stop codons were replaced with coding sequences of *ScURA3* and *ScHIS3* as described (70). Knockout mutants were authenticated by PCR using external primers (Table S4). Cells were cultured in synthetic complete 2% dextrose (SCD) medium at 30° C.

### Genome-wide screens

Large pools of Hermes transposon insertion mutants in strains BG14 (wild-type) and CGM1094 (*pdr1Δ::HYGr*) were thawed from frozen stocks (28), grown to stationary phase in synthetic complete 2% dextrose (SCD) medium, diluted 100-fold into fresh medium containing or lacking 1 µg/mL FK506 (SelleckChem), and shaken for one day at 30°C. Cells were then pelleted, washed once in SCD medium, resuspended in an equal volume of fresh SCD medium, and shaken for one day at 30°C. Cells were then pelleted, resuspended in 30 mL of 15% glycerol, and frozen in aliquots at −80°C. Genomic DNA was extracted from the aliquots, sheared by sonication, A-tailed, ligated to splinkerette adapters, PCR amplified, and sequenced using a MiSeq (Illumina) instrument as described previously (30). Sequence reads were demultiplexed, mapped to the BG2 reference genome, filtered for quality, and then tabulated gene-wise (30). The tabulated data were normalized, and then a z-score was calculated for each gene using the log_2_ ratio of transposon insertions in FK506 versus control divided by the local standard deviation, which was estimated from the data as described previously (30).

### Spot tests of drug susceptibility

Single colonies were picked and grown to saturation in SCD medium, serially diluted in 5-fold increments, and frogged to agar plates containing SCD medium with or without supplements of FK506 (1 µg/mL) or cyclosporin A (100 µg/mL; SelleckChem). Strains were grown at 30°C for 24 hours. Images were taken on a Gel Doc XR+ (Image Lab, BioRad).

### RT-PCR experiments

Single colonies were picked and grown overnight at 30°C to mid-log phase in SCD medium. For each sample, cells were diluted to OD600 = 0.1 in fresh SCD medium with or without the stressor (see below) and shaken at 30°C. At the appropriate time points, 1.5 mL of the culture was harvested by centrifugation (14 k, 60 sec) and the supernatant aspirated. Cell pellets were flash frozen in liquid nitrogen and stored at −80°C until RNA extraction. Total RNA was extracted from cells using a hot acid phenol-chloroform extraction protocol (71). Briefly, cells were lysed in an RNA lysis buffer (6 mM NaOAc, 8.4 mM EDTA, 1% SDS), and then RNA was purified through two phenol extractions and a final chloroform extraction. RNA was precipitated with isopropanol and resuspended in TE buffer. RNA extracts were treated with DNAse (New England Biolabs) to ensure no genomic DNA contamination. 1 µg of RNA was reverse transcribed using the High-Capacity cDNA Reverse Transcription Kit (Thermo). Real-time PCR was performed using the CFX96 Touch Real-Time PCR Detection System (BioRad) using the ABsolute Blue QPCR Mix SYBR Green Kit (ThermoFisher) with the following parameters: 15 min at 95°C, 40x (15 min at 95°C, 30 min at 58°C, 30 min at 72°C). Target gene transcript levels were normalized to averaged *TEF1* and *PGK1* transcript levels in each sample, and this ratio from each sample was normalized to that of untreated BG14 cells. Target primers are identified in Table S4.

### Cell Death assay

Single colonies were picked and grown to log phase at 30°C in SCD medium. Cells were back-diluted to an OD600 = 0.1 and dosed with 0.6 µg/mL manogepix (SeleckChem), 20 µg/mL tunicamycin (Tocris Bioscience), or 0.12 µg/mL micafungin (Cayman Chemicals) and continued to grow at 30° C. Samples were taken at time-points, spun down, stained with propidium iodide (100 µg/mL) in PBS, and manually counted on a fluorescence microscope (Zeiss Axioscope). PI-positive cells were tallied out of 200 cells counted per sample.

## ACKNOWLEDGMENTS

The authors thank Drs. Brendan Cormack and Alejandro de las Peñas for generously providing *C. glabrata* strains and advice. We are grateful to Drs. Winston Timp, John Kim, and Andrew Gordus for providing access to critical instruments. Drs. Lars Essen and Hans-Ulrich Mosch provided helpful advice and insights. Josh Schultz commented thoughtfully on the project and the manuscript. This research was supported by grants from the National Institutes of Health (T32-GM007231 to the JHU CMDB training program; R01-AI153414 to KWC).

